# A Trifector of New Insights into Ovine Footrot for Infection Drivers, Immune Response and Host Pathogen Interactions

**DOI:** 10.1101/2021.05.13.444115

**Authors:** Adam M. Blanchard, Ceri E. Staley, Laurence Shaw, Sean R Wattegedera, Christina-Marie Baumbach, Jule K. Michler, Catrin Rutland, Charlotte Back, Nerissa Newbold, Gary Entrican, Sabine Tötemeyer

**Author notes:** Bristol Veterinary School, University of Bristol, Langford House, Langford, Somerset, BS40 5DU (Nerissa Newbold). The Roslin Institute, The University of Edinburgh, Easter Bush, Scotland, EH25 9RG (Gary Entrican).

## Abstract

Footrot is a polymicrobial infectious disease in sheep causing severe lameness, leading to one of the industry’s biggest welfare problems. The complex aetiology of footrot makes in-situ or in-vitro investigations difficult. Computational methods offer a solution to understanding the bacteria involved, how they may interact with the host and ultimately providing a way to identify targets for future hypotheses driven investigative work. Here we present the first combined global analysis of the bacterial community transcripts together with the host immune response in healthy and diseased ovine feet during a natural polymicrobial infection state using metatranscriptomics. The intra tissue and surface bacterial populations and the most abundant bacterial transcriptome were analysed, demonstrating footrot affected skin has a reduced diversity and increased abundances of, not only the causative bacteria *Dichelobacter nodosus*, but other species such as *Mycoplasma fermentans* and *Porphyromonas asaccharolytica*. Host transcriptomics reveals a suppression of biological processes relating to skin barrier function, vascular functions, and immunosurveillance in unhealthy interdigital skin, supported by histological findings that type I collagen (associated with scar tissue formation) is significantly increased in footrot affected interdigital skin comparted to outwardly healthy skin. Finally, we provide some interesting indications of host and pathogen interactions associated with virulence genes and the host spliceosome which could lead to the identification of future therapeutic targets.

**Impact Statement:** Lameness in sheep is a global welfare and economic concern and footrot is the leading cause of lameness, affecting up to 70% of flocks in the U.K. Current methods for control of this disease are labour intensive and account for approximately 65% of antibiotic use in sheep farming, whilst preventative vaccines suffer from poor efficacy due to antigen competition. Our limited understanding of cofounders, such as strain variation and polymicrobial nature of infection mean new efficacious, affordable and scalable control measures are not receiving much attention. Here we examine the surface and intracellular bacterial populations and propose potential interactions with the host. Identification of these key bacterial species involved in the initiation and progression of disease and the host immune mechanisms could help form the basis of new therapies.

## Introduction

Ovine footrot is a persistent animal welfare issue and has a significant financial burden for farmers due to the cost of preventative footbaths, antibiotic treatments, and reduced carcass weights at slaughter (1). The causative bacterium *Dichelobacter nodosus* (*D. nodosus*) has received extensive attention since its description in the initiation of footrot (2). However, it has been accepted since the beginning of the 20^th^ century that footrot is a polymicrobial disease, with *Fusobacterium necrophorum (F. necrophorum)*, *Spirochaeta penortha (S. penortha)* (3), *Treponema podovis (T. podovis)* (4) and *Corynebacterium pyogenes (C. pyogenes)* (5) proposed as species that can exacerbate the lesions.

Currently our understanding of bacterial populations associated with footrot is only based on 16S rRNA analysis from the skin surface (6, 7). The highly abundant genera identified in the footrot samples were congruent with those identified previously by standard microbiological techniques (*Corynebacterium, Fusobacterium, Dichelobacter* and *Treponema*). However, additional genera were also identified (*Mycoplasma, Psychrobacter* and *Porphyromonas*) (7) and their absence using traditional culture techniques, could be due to the fastidious nature of the bacteria (8) or that they were not yet identified (9). Investigating the total bacterial load within tissues, we have shown recently, that in healthy tissues, bacterial load is similar throughout tissue depth and did not extend beyond the follicular depth in the reticular dermis. In contrast, in footrot samples, the bacterial load was highest in the superficial (or cornified) epidermal layers and decreasing in the deeper layers but still beyond follicular depth (10). This suggests that the infection allows for further invasion from other species of bacteria to penetrate deeper into the interdigital tissue, however these data were limited to presence of bacteria based on universal primers not allowing to identify species.

There is also a lack of information regarding the host infection, how an immune response is mounted and the species interactions. This area of investigation has recently benefitted from the use of metatranscriptomics, a method of assessing host-pathogen interactions based on associated gene expression changes (11). The use of metatranscriptomics has been reviewed extensively (12), however, current published methods are based on, or optimisations of, cell culture models as developed in the original methods article (11). The use of metatranscriptomics in natural polymicrobial infections is not as well reported. The first documented use was in relation to the onset of paediatric asthma (13), and oral disease (14, 15). However, its application to Bovine Digital Dermatitis (BDD) (16), which has a similar clinical presentation and bacteria associated with footrot (17–20) highlighted its suitability to further our understanding of the inter-cellular microbial populations associated with agricultural diseases. Utilising this experimental design, we have been able to determine the bacterial populations on the surface of the interdigital skin and within the deeper infected tissue, identify the differential expression of the host transcripts and elucidate interactions between the host and bacteria.

## Results

### Sequence data

Foot swabs and whole thickness skin biopsies were collected from sheep post slaughter that had at least one apparently healthy foot (n=13) and one with signs of footrot (n=13) to obtain matching samples from the same sheep. After quality filtering there was an average of 8.7 million discordant ovine reads per sample to be used for bacterial taxonomic assignment from the foot biopsies, and 20.8 million discordant ovine reads for the accompanying swabs. All reads had an average phred score of 40. Diversity statistics were calculated for each sample after assignment. Using the Shannon indices and calculating an equitability score (natural log of the species richness) representing a maximum diversity, revealed that healthy feet were highly diverse but footrot feet showed a reduction in diversity (Table 1, Fig 1A). The Simpson indices also indicated that there was more diversity in the healthy samples with an average of 0.78 in healthy compared to 0.69 in the footrot biopsies. (Figure 1B). This was furthermore reflected in the swabs with an average of 0.94 in the healthy samples compared to 0.77 in footrot (Figure 1). The Shannon index showed a significant difference between the two conditions for both, biopsies (p=≤0.005) and swabs (p=≤0.05), whereas only the swabs showed a significant difference for the Simpson index (p=≤0.005; Figure 1).

**Figure 1.**
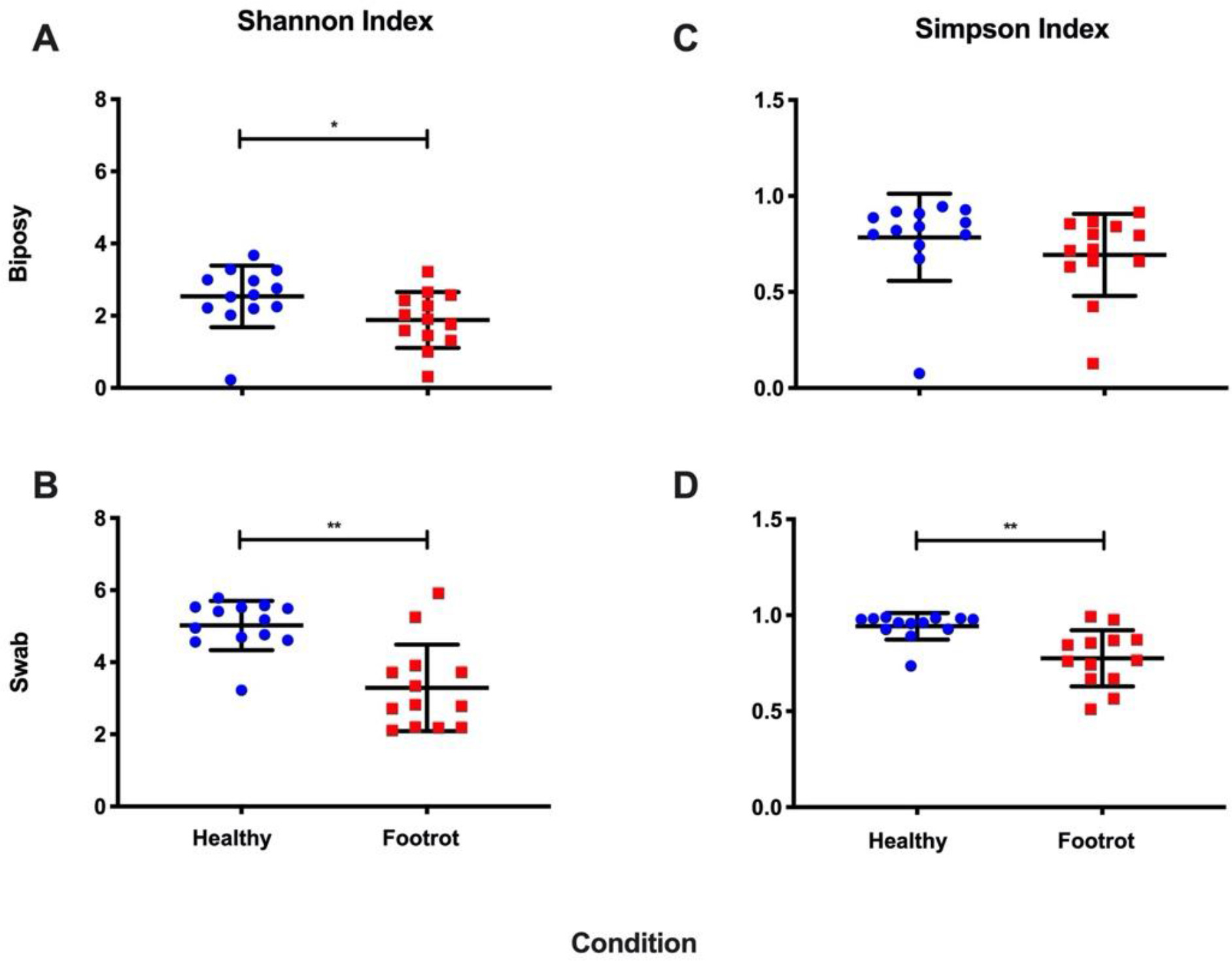
Diversity statistics for the biopsy and swab samples. A) Shannon Index of biopsy samples, B) Shannon Index of swab samples, C) Simpson Index of biopsy samples, D) Simpson index of swab samples. Significant decreases were observed from footrot affected samples, for swabs using both Shannon and Simpson indices. A significant decrease in footrot affected samples was only observed for biopsies using the Shannon index. Statistical significance calculated using Mann Whitney U (* p=0.05, ** p=0.005, *** p=0.0005).

**Figure 2.**
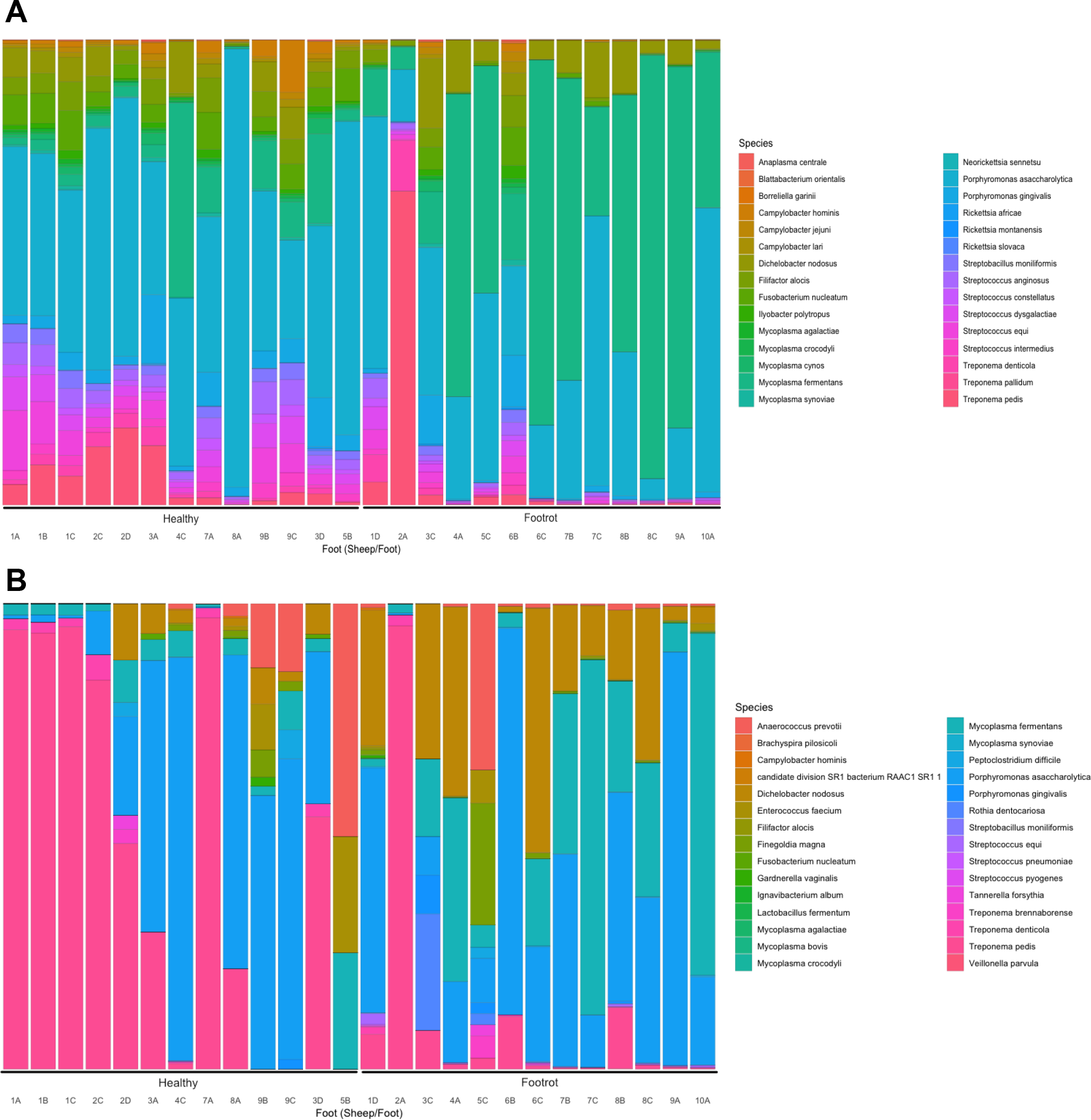
Species of bacteria identified as increased in abundance in footrot affected feet when compared to healthy feet. A) Shows the top 30 species of bacteria in swab samples and B) Shows the top 30 species of bacteria in biopsy samples.

**Table 1.** Comparison of average calculated and maximum diversity for each condition. Demonstrating the overall reduction in bacterial community diversity for footrot affected individuals when compared to the calculated maximum diversity expected from the data.

| Condition | Biopsy<br>Equitability | Biopsy<br>Shannon | Swab<br>Equitability | Swab<br>Shannon |
| --- | --- | --- | --- | --- |
| Healthy | 3.9 | 2.5 | 7.1 | 5.0 |
| Footrot | 3.3 | 1.9 | 7.1 | 3.2 |

### Bacterial community

Differences in abundance calculated between the two conditions were identified as samples having a >2 log fold change, with an FDR (Benjamini-Hochberg) corrected p-value <0.05 and where average counts had a difference greater than 10 (full taxonomic assignments are available in Supplementary Table 1 for swabs and Supplementary Table 2 for biopsies). In swabs, 20 species of bacteria were found in significantly increased abundance in footrot samples. These included *T. pedis, T. denticola, D. nodosus*, and *F. necrophorum,* all known to cause various foot diseases in sheep. Among the bacterial species found in significantly reduced abundance in footrot samples were ten species of *Staphylococcus* spp., *Bacillus licheniformis*, *Parageobacillus thermoglucosidasius* and *Nocardiopsis alba*. All differential abundance data for the swabs are available in Supplementary Table 3.

Applying the same criteria to biopsies, three species of bacteria were found in differential abundance between the two conditions, namely *D. nodosus, Mycoplasma fermentans (M. fermentans)* and *Porphyromonas asaccharolytica (P. asaccharolytica). D. nodosus* had the most significant increase in footrot biopsies with a log-fold change increase of 7.0 (p=1.89E-06), *M. fermentans* had a log-fold change of 6.2 (p=2.59E-05) whilst *P. asaccharolytica* had a log-fold increase of 3.5 (p=0.018). No species were found to be significantly decreased between the two conditions in the biopsies (all differential abundance data for the biopsies are available in Supplementary Table 4). Although some archaea were identified in both biopsy and swab samples, none were significantly more or less abundant in footrot affected feet compared to healthy feet.

As short read sequencing has limitations in identifying bacteria to species level, the most significantly increased abundant bacteria in footrot affected tissues were confirmed to be *D. nodosus*, *M. fermentans* and *P. asaccharolytica,* by specific qPCR, species specific PCR and PCR followed by sequencing, respectively. In addition, *F. nucleatum* was identified as the only *Fusobacterium* species, however qPCR demonstrated this as misidentification and *F. necrophorum* was present as expected.

### Comparative analysis of in tissue and surface bacterial communities

The taxonomic assignments from both, swab and biopsy data, were tested to ascertain whether a clear relationship existed between taxonomic assignments for the same sheep using both the correlation and similarity hypothesis tests outlined in the methods section. Under the null hypothesis for the correlation test, there was no correlation between swab and tissue samples. A p-value of 0.0301 was obtained, providing strong evidence of a relationship. However, it should be noted that this is evidence of a relationship in the presence of bacterial species between biopsy and swab samples rather than them containing the same species.

To test the latter claim, the similarity test from the methods section was used. Here, the null hypothesis was that biopsy and swab samples reveal the presence of the same bacteria. This test produced a conservative p-value of < 10^-5, providing over overwhelming evidence that swab and biopsy samples from the same sheep do not contain the same species of bacteria. Specifically, two random biopsy samples will have more species in common than a swab and biopsy from the same sheep.

### Differential expression of pro-inflammatory mediators in healthy versus footrot affected interdigital skin

Among the transcripts that showed increased expression in footrot-affected interdigital skin were a large number of proteins important for barrier function. These included proteins involved in collagen production and collagen binding (Procollagen C-endopeptidase enhancer 2 [PCOLCE2], Collagen Type VI alpha 6 chain, Collagen Type XXIII alpha 1 chain and keratocan/lumican [collagen-binding leucine-rich proteoglycans widely distributed in interstitial connective tissues]; cell-cell adhesion (cadherin[CDH]3, CDH19,pro[P]CDH10); maintenance of cell junctions (GJB4) and long chain fatty acid synthesis (fatty acid elongase [ELOVL]7, ELOVL3, acyl-CoA synthetase bubblegum family member [ACSBG]1), and acyl-CoA wax alcohol acyltransferase [AWAT]1). In addition, transcripts involved in immunosurveillance such as scavenger receptors SCARA5 and SSC5D were more highly expressed in footrot affected samples (see Table 2 for top 25 transcripts, Supplementary Table 5 for all transcripts). In contrast, transcripts that showed lower expression compared to healthy interdigital skin include cytokines involved in wound-healing (IL-19, IL-20) and keratinocyte proliferation/differentiation (IL-6 and leukaemia inhibitory factor [LIF]), epithelial cell-derived chemokines that recruit monocytes (CCL2), lymphocytes (CCL20) and neutrophils (CXCL1, CXCL8) and prostaglandin-endoperoxide synthase 2 (PGE2/COX2), which is also involved in skin wound healing. Another group of significantly decreased transcripts include matrix metalloproteases (MMP1, MMP3, MMP9, MMP13, MMP20, (tenacin C) TNC, TIMP1) and their regulators (SERPINE1, ADAMTS4, ADAMTS16) associated with chronic wounds and collagen turnover (see Table 3 for top 25 transcripts, Supplementary Table 5 for all transcripts).

**Table 2:** Top 25 differentially higher expressed genes in footrot affected skin when compared to healthy skin.

| Gene name | Description | β-value | Standard error | q-value |
| --- | --- | --- | --- | --- |
| ELOVL7 | ELOVL fatty acid elongase 7 | 2.812237 | 0.884427 | 0.035516 |
| MNT | MAX network transcriptional repressor | 2.415885 | 0.773256 | 0.038156 |
| FRAS1 | Fraser extracellular matrix complex subunit 1 | 2.358302 | 0.738887 | 0.034774 |
| CA6 | carbonic anhydrase 6 | 2.273248 | 0.686033 | 0.029014 |
| ATP13A4 | ATPase 13A4 | 2.261712 | 0.648362 | 0.023149 |
| PNPLA5 | patatin like phospholipase domain containing 5 | 2.230351 | 0.627058 | 0.021215 |
| ELOVL3 | ELOVL fatty acid elongase 3 | 2.070317 | 0.598268 | 0.02425 |
| STMN2 | stathmin 2 | 2.021979 | 0.464736 | 0.007856 |
| NOS1 | nitric oxide synthase 1 | 2.004654 | 0.546986 | 0.018297 |
| ACSBG1 | acyl-CoA synthetase bubblegum family member 1 | 1.933404 | 0.590342 | 0.030742 |
| AWAT1 | acyl-CoA wax alcohol acyltransferase 1 | 1.849357 | 0.543633 | 0.02594 |
| CCL26 | C-C motif chemokine ligand 26 | 1.842727 | 0.621591 | 0.046882 |
| CCDC155 | coiled-coil domain containing 155 | 1.791928 | 0.584831 | 0.041351 |
| PI16 | peptidase inhibitor 16 | 1.765247 | 0.400402 | 0.007477 |
| FAR2 | fatty acyl-CoA reductase 2 | 1.711846 | 0.519782 | 0.02993 |
| PTX4 | pentraxin 4 | 1.681009 | 0.336588 | 0.004979 |
| LRRC36 | leucine rich repeat containing 36 | 1.669794 | 0.465845 | 0.020504 |
| AGTR1 | angiotensin II receptor type 1 | 1.661825 | 0.442673 | 0.016379 |
| GALNT8 | polypeptide N-acetylgalactosaminyltransferase 8 | 1.631845 | 0.457915 | 0.020974 |
| CYP2F1 | cytochrome P450 family 2 subfamily F member 1 | 1.55567 | 0.44436 | 0.022765 |
| TOGARAM2 | TOG array regulator of axonemal microtubules 2 | 1.530427 | 0.48021 | 0.035066 |
| DNASE1L2 | deoxyribonuclease 1 like 2 | 1.515543 | 0.406144 | 0.0169 |
| FAM221A | family with sequence similarity 221 member A | 1.506933 | 0.378682 | 0.01257 |
| AQP9 | aquaporin 9 | 1.506146 | 0.384521 | 0.013256 |
| DGAT2L6 | diacylglycerol O-acyltransferase 2 like 6 | 1.505104 | 0.480771 | 0.03797 |

qval: FDR adjusted p-value using Benjamini-Hochberg; $\beta$ -value: bias estimator analogous to fold change

**Table 3:** Top 25 lower expressed genes in footrot affected interdigital skin compared to healthy skin.

| Gene name | Description | β-value | Standard error | q-value |
| --- | --- | --- | --- | --- |
| IL-19 | interleukin 19 | -2.96775 | 0.804409 | 0.017781 |
| PTGS2 | prostaglandin-endoperoxide synthase 2 (PGE2/COX2) | -2.8683 | 0.75813 | 0.015736 |
| MMP3 | matrix metalloproteinase 3 | -2.71059 | 0.616087 | 0.007597 |
| IL-6 | interleukin-6 precursor | -2.4267 | 0.568937 | 0.008635 |
| IL-20 | interleukin 20 | -2.30205 | 0.758643 | 0.04305 |
| A2ML1 | alpha-2-macroglobulin like 1 | -2.23879 | 0.765254 | 0.049443 |
| CCL20 | C-C motif chemokine ligand 20 | -2.18479 | 0.501858 | 0.007856 |
| CXCL8 | Interleukin-8 | -2.10659 | 0.685411 | 0.040981 |
| SLPI | antileukoproteinase precursor | -2.04964 | 0.672506 | 0.042275 |
| MGAM | maltase-glucoamylase | -2.02022 | 0.568845 | 0.021334 |
| MARCKSL1 | MARCKS like 1 | -1.95955 | 0.631205 | 0.039119 |
| MEFV | MEFV, pyrin innate immunity regulator | -1.90552 | 0.514121 | 0.017474 |
| MMP13* | matrix metalloproteinase 13 | -1.83676 | 0.523263 | 0.022523 |
| ADAMTS16 | ADAM metalloproteinase with thrombospondin type 1 motif 16 | -1.80019 | 0.479709 | 0.016379 |
| ACOD1 | aconitate decarboxylase 1 | -1.68652 | 0.563289 | 0.045291 |
| PTX3 | pentraxin 3 | -1.66311 | 0.446328 | 0.016904 |
| MMP13* | matrix metalloproteinase 13 | -1.65481 | 0.561768 | 0.048097 |
| ADAMTS4 | ADAM metalloproteinase with thrombospondin type 1 motif 4 | -1.59355 | 0.394111 | 0.011418 |
| MMP1 | matrix metalloproteinase 1 | -1.55518 | 0.37913 | 0.01037 |
| FOSL1 | FOS like 1, AP-1 transcription factor subunit | -1.51685 | 0.383897 | 0.012932 |
| MGAT3 | mannosyl (beta-1,4-)-glycoprotein beta-1,4-N-acetylglucosaminyltransferase | -1.46074 | 0.377507 | 0.014131 |
| CCL2 | C-C motif chemokine ligand 2 | -1.40425 | 0.325691 | 0.008232 |
| CSF3 | colony stimulating factor 3 | -1.4015 | 0.473053 | 0.047061 |
| PLAUR | urokinase plasminogen activator surface receptor precursor | -1.36363 | 0.322566 | 0.009128 |

qval: FDR adjusted p-value using Benjamini-Hochberg; $\beta$ -value: bias estimator analogous to fold change
\*Splice variants

### Biological process enrichment

Using BCCC biclustering and associated GO biological processes, the genes and conditions grouped into a total of 32 clusters. There were 2531 genes over 20 samples that clustered showing upregulated biological processes including significant positive regulation. The top 15 were positive regulation of transcription (n=106, p=2.94e-26), protein folding (n=34, p=3.5e-21), regulation of DNA templated transcription (n=121, p=4.51e-21), metabolic process (n=61, p=1.44e-18), DNA templated transcription (n=63, p=1.48-18), rRNA processing (n=20, p=9.12e-15), protein transport (n=32, p=2.08e-13), ribosome biogenesis (n=17, p=2.20e-13), osteoblast differentiation (n=24, p=4.01e-12), transcription from RNA polymerase II promotor (n=49, p=5.84e-12), negative regulation of transcript from RNA polymerase II promotor (n=63, p=7.75e-12), negative regulation of apoptotic process (n=46, p=1.91e-11), positive regulation of telomerase RNA localisation to cajal body (n=10, p9.34e-11), proteolysis (n=63, p=1.65e-10) and translational initiation (n=16, p=1.74e-10).

There were 541 genes over 17 samples that clustered showing significant downregulation of biological process in footrot affected samples. Cluster one showed a decrease in epidermis development (n=17, p=2.3e-06), multicellular organismal water homeostasis (n=8, p=1.8e-05), peptidoglycan catabolic processes (n=4, p=4.6e-05), antimicrobial humoral response (n=7, p=9.6e-05), tissue development (n=41, p=1.1e-04), monovalent inorganic cation homeostasis (n=8, p=8.2e-4), defence response to bacteria (n=11, p=8.2e-04), fatty acid metabolic processes (n=12, p=1.3e-03), polyol transport (n=3, p=2.7e-03), skin development (n=11, p=2.7e-03), water transport (n=4, p=3.4e-03) and regulation of pH (n=6, p=3.4e-03). Cluster two showed downregulation for neutrophil chemotaxis (n=7, p=4.3-e06), myeloid leukocyte migration (n=9, p=7.2e-06), leukocyte migration (n=11, p=9.8e-06), cell chemotaxis (n=10, p=2.6e-05), defence response (n=19, p=2.8e-05), immune system processes (n=26, p=4.3e-05), chemotaxis (n=12, p=1.2e-04), antimicrobial humoral response (n=5, p=1.2e-04), immune response (n=18, p=1.2e-04) and response to external stimulus (n=23, p=1.5e-04). The third cluster showed downregulation for S-adenosylhomocysteine catabolic process (n=2, p=5.7e-04) alone.

Investigating KEGG pathway enrichment also identified cytokine-cytokine receptor interaction (n=26, p=4.5e-6), IL-17 signalling pathway (n=13, p=7.7e-08), TNF signalling pathway (n=12, p=7.7e-05) to be downregulated in the footrot affected samples. Whilst steroid hormone biosynthesis (n=7, p=2.4e-04) and ribosome biogenesis (n=31, p=4.07e-08) were upregulated.

### Putative Host pathogen interactions

The bacterial RNA reads were aligned against the bacterial transcriptomes that were identified as those traditionally associated with ovine foot disease (*D. nodosus, F. necrophorum*, *T. pedis* and *T. denticola*) and those additionally found to be the most differentially abundant in the footrot biopsy samples *(M. fermentans* and *P. asaccharolytica*). These data were then used, along with the host expression data to understand correlations and host and pathogen interactions (Supplementary Table 6).

The interactions were calculated with an FDR adjusted p-value using the Benjamini-Hochberg procedure. Due to the stringency of this multiple test adjustment, no significance was determined. However, the raw p-values were low (in some cases < 0.00005), therefore these data were investigated further but only as an indicative positive bacteria/gene correlation. Based on a raw p-value of <0.00005 there were four sheep transcripts that were associated with five *D. nodosus* genes (Table 4). From *D. nodosus* aminoacyl-histidine dipeptidase, acidic extracellular subtilisin-like protease precursor (AprV5), outer membrane protein 1E, Bacterial extracellular solute-binding protein and aminoacyl-histidine dipeptidase were identified to correlate with small nucleolar RNA, C/D box, U6 spliceosomal RNA, synapsin and U6 spliceosomal RNA from *Ovis aries*. There were more correlations between *M. fermentans* and *Ovis aries* with a total of 15 bacterial transcripts associated with four host transcripts where raw p=0.0005. The bacterial transcripts were shown to be overwhelmingly responsible for cellular transport on both the host and pathogen side. There were a further three bacterial and sheep interactions in *T. pedis* (p=0.00005) which suggested the bacterial flagellin and host membrane protein, and a bacterial hypothetical protein and putative lipoprotein correlates with a sheep miRNA (Table 4).

**Table 4.** Correlations between bacterial and *host* gene expression.

| Bacterial Species | Protein Accession | Function | *Ovis aries* Transcript accession | Function | p-value |
| --- | --- | --- | --- | --- | --- |
| D. nodosus | ABQ13122.1 | aminoacyl-histidine dipeptidase | ENSOART00000026163 | small nucleolar RNA, C/D box | 1.66E-05 |
| D. nodosus | ABQ13667.1 | acidic extracellular subtilisin-like protease precursor (AprV5) | ENSOART00000023789 | U6 spliceosomal RNA | 7.10E-05 |
| D. nodosus | ABQ13351.1 | outer membrane protein 1E | ENSOART00000014155 | synapsin | 7.49E-05 |
| D. nodosus | ABQ13881.1 | Bacterial extracellular solute-binding protein |  |  | 7.49E-05 |
| D. nodosus | ABQ13122.1 | aminoacyl-histidine dipeptidase | ENSOART00000023789 | U6 spliceosomal RNA | 7.49E-05 |
| M. fermentans | WP_013526775.1 | Protein translocase subunit | ENSOART00000015973 | Pro-platelet basic protein | 2.88E-05 |
| M. fermentans | WP_013526734.1 | Putative Oligopeptide ABC transporter, ATP-binding protein | ENSOART00000022767 | Novel Transcript | 4.71E-05 |
| M. fermentans | WP_013526734.1 | DNA-directed RNA polymerase subunit beta | ENSOART00000013696 | Potassium voltage-gated channel | 7.04E-05 |
| M. fermentans | WP_013526778.1 | ATP synthase subunit beta | ENSOART00000017736 | Novel Transcript | 7.04E-05 |
| M. fermentans | ADV34079.1 | MgpA like protein | ENSOART00000013889 | CXXC Type Zinc Finger 1 CPG Binding PHD Finger | 9.28E-05 |
| M. fermentans | WP_013354336.1 | bifunctional oligoribonuclease/PAP phosphatase |  |  | 9.28E-05 |
| M. fermentans | WP_013526633.1 | Adenine phosphoribosyltransferase |  |  | 9.28E-05 |
| M. fermentans | ADV34286.1 | Oligopeptide ABC transporter permease protein |  |  | 9.28E-05 |
| M. fermentans | WP_013354483.1 | ABC transporter permease |  |  | 9.28E-05 |
| M. fermentans | WP_013354556.1 | sugar ABC transporter permease |  |  | 9.28E-05 |
| M. fermentans | ADV34629.1 | NADPH flavin oxidoreductase |  |  | 9.28E-05 |
| M. fermentans | WP_013354747.1 | nitroreductase family protein |  |  | 9.28E-05 |
| M. fermentans | ADV34954.1 | Transcription antitermination protein |  |  | 9.28E-05 |
| M. fermentans | WP_013527166.1 | ABC transporter ATP-binding protein |  |  | 9.28E-05 |
| M. fermentans | WP_013527168.1 | ABC transporter permease |  |  | 2.88E-05 |
| T. pedis | WP_024465740.1 | Flagellin | ENSOART00000022912 | Novel Membrane Protein | 5.38E-05 |
| T. pedis | WP_051150643.1 | Hypothetical Protein | ENSOART00000026139 | ncRNA (MiRNA) | 5.47E-05 |
| T. pedis | AGT42887.1 | Putative Lipoprotein |  |  | 5.47E-05 |

The correlations between the sheep transcripts and the bacterial transcripts from *F. necrophorum*, *P. asaccharolytica* and *T. denticola* had raw p-values of <0.0005, <0.001 and <0.004, respectively. Although low, p-values with the number of tests being performed they were not investigated any further (full data is available in Supplementary Table 6).

### Collagen composition differs in the dermis of healthy and footrot tissues

Since collagen composition changes in scar tissue formation, picrosirius stained tissue sections were used to differentiate collagen type I and III from each other and other collagen types. To investigate whether there were any differences in the collagen composition of the dermis, the proportions of type I, type III, non-differentiated and total collagen were calculated (Fig 4). The proportions of total and type I collagen were significantly increased in the dermis of footrot samples compared to healthy samples (p=0.04 and 0.042, respectively, Fig 4 A, D). The proportions of non-differentiated and type III collagen were not significantly different in the dermis layers of healthy and footrot-affected tissues (Fig 4, B-C).

**Figure 4:**
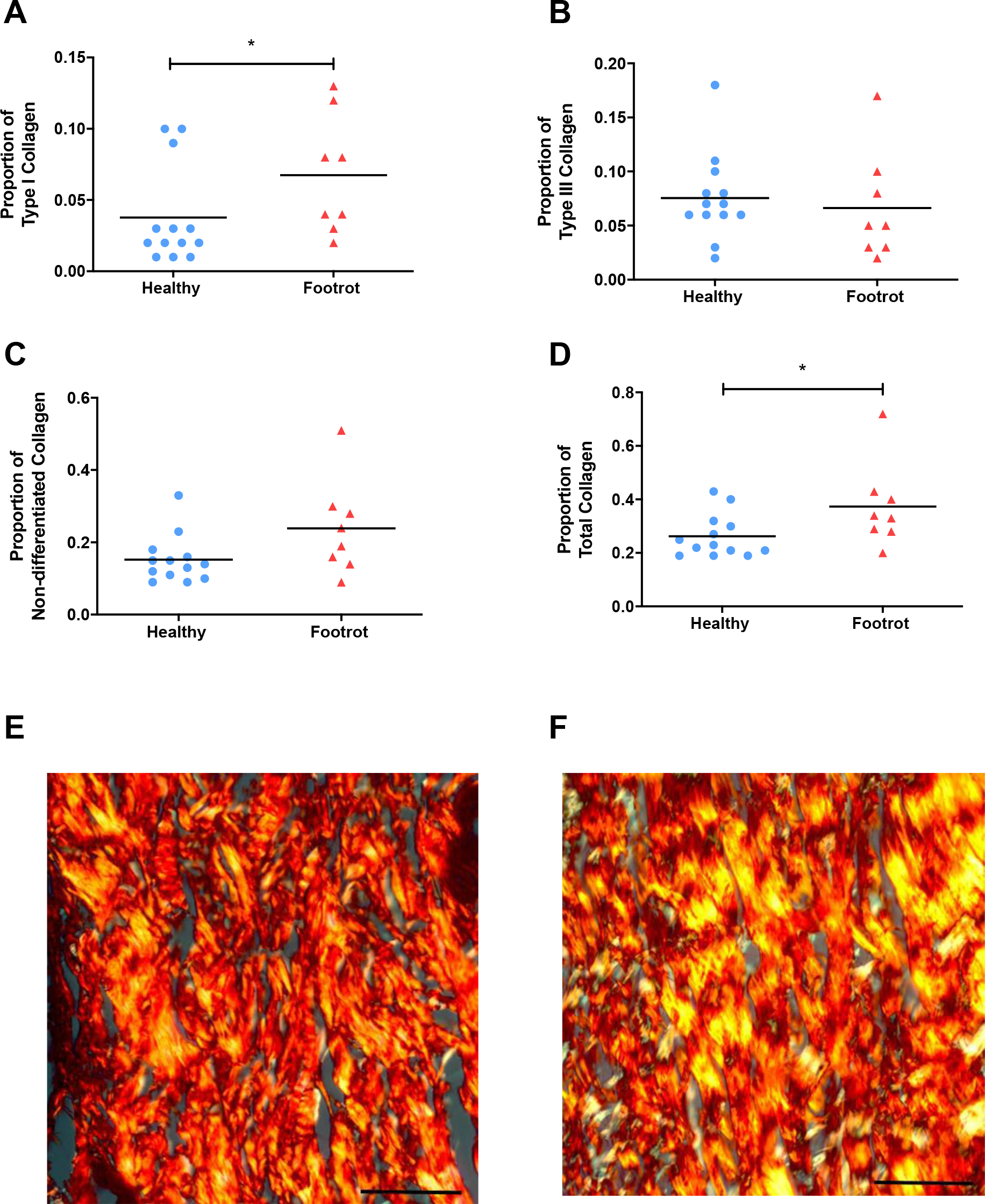
Collagen expression in healthy and footrot ovine interdigital skin dermis. Picrosirius histological staining was used to differentiate and quantify collagens in healthy (n=13) and footrot (n=8) samples. Proportions of type I collagen (A), type III collagen (B), undifferentiated collagens (C) and total proportion of collagen (D). Representative photomicrographs showing picrosirius staining under phase microscopy from healthy (E) and footrot (F) samples, type I collagen stained yellow, type III collagen stained green, undifferentiated collagen stained red. Scale bars represent 50µm. Significance is designated by asterisk on a straight line with the T-test result defined as * = *p*≤0.05.

## Discussion

*D. nodosus,* was established as the causative bacterium of ovine footrot in the 1940’s, and it has long been accepted that *F. necrophorum* plays a role in the disease aetiology (4). However, in our work we have identified additional common core species that are also associated with footrot lesions (*M. fermentans* and *P. asaccharolytica*) and we have shown putative interactions with the molecular host defence systems during infection in bacterial species identified as highly abundant in footrot and even without a significant difference in abundance between healthy and footrot affected such as *T. pedis*. Using paired biopsies collected from footrot affected sheep at point of slaughter we were able to comprehensively show that skin swabs are a poor proxy for identifying what bacteria are present in the tissue. This is potentially due to the bacterial contamination present from environmental sources such as faeces and soil collected during transport and grazing. However, we have shown that biopsies provide an intradermal approach to reproducibly assess differences between individual animals in an invasive infection like footrot.

### The bacterial community structure of footrot

Previous studies have shown the ovine interdigital bacterial community structure using 16S rDNA, with predominant bacterial genera identified as *Mycoplasma spp, Corynebacterium spp, Psychobacter spp, Treponema spp, Staphylococcus spp, Peptostreptococcus spp* and *Dichelobacter spp* (6, 7). The results from this study are highly congruent with what has previously been identified, however due to the greater taxonomic sensitivity afforded by metagenomics and metatranscriptomics we have been able to classify those bacterial genera to a species level. Those with differential abundance associated with footrot found on the skin surface were identified as *T. pedis, T. denticola, D. nodosus*, and *F. necrophorum.* These species are commonly found with other ovine foot diseases, contagious ovine digital dermatitis (CODD) (21) and interdigital dermatitis (ID) (7) and the bovine foot disease bovine digital dermatitis (BDD) (18). The species with differential abundance associated with footrot intradermally were *D. nodosus, M. fermentans* and *P. asaccharolytica.* Given that *D. nodosus* is a poor pathogen and often requires tissue damage and the presence of other bacteria to infect, it stands to assume *P. asaccharolytica and M. fermentans* may also have an important role in disease susceptibility. These differences between healthy and footrot affected feet also extended beyond presence and absence of species to the overall bacterial diversity, with a significant drop in footrot samples. This reduction in diversity has been mirrored in CODD (21).

The benefits of using metagenomics compared to 16S rDNA studies include that the scope can be extended to incorporate archaeal and DNA virus discovery. In the current study there was a lack of correlation between either and the disease state, however, this may be unsurprising without an enrichment step or optimal DNA extraction to make them specifically more suitable for viral or archaeal identification, resulting in a poor representation for those species identified.

### Host response and pathogen interactions

Investigating the host pathogen interactions through correlation analysis has identified some interesting associations which warrant further investigations. The most promising appears to be the association between virulence gene *aprV5* and the *Ovis aries* transcript U6 spliceosomal RNA. This particular non-coding small nuclear RNA (snRNA) is responsible for catalysing the excision of introns and is a major aspect of post translation modifications, with the ability to alter the structure, function and stability of the translated protein. In the case of infections, some species of bacteria have been implicated in hijacking the host splicing machinery and altering the splicing pattern leading to the perturbation of the host response (22, 23). Despite the lack of knowledge around the mechanism, there is evidence that certain *Listeria, Salmonella* and *Mycobacterium* species have the ability to produce factors that have a direct or indirect impact on the regulation of alternative splicing (23–25). Alternative splicing from the U6 spliceosomal RNA can interfere with the normal activation of T cell and B lymphocytes and the regulation of the signalling in several TLR’s (TLR2, TLR3 and TLR4) (26), which could tie in with certain pathways (monovalent inorganic cation homeostasis, defence response to bacteria, skin development, neutrophil chemotaxis, leukocyte migration, defence response, immune system processes, immune response) which were identified as being downregulated in these data.

The acidic extracellular protease *aprV5 is* associated with the correct cleavage of the other proteases secreted by *D. nodosus,* AprV2 and BprV, to their mature active form (Han et al., 2012). Whereas the closely related AprV2 acidic protease is a known virulence factor responsible for elastase activity and its degradation of the host extra cellular matrix (28), the role of AprV5 in footrot is unclear. The abundance of isolates with *aprV5* has been shown to be around 25% from clinically affected farms and lacks any clear delineation between disease severity (29). However, as viral-mediated proteases have been implicated in the degradation of host small non-coding ribonucleal proteins (snRNP) (30) and other bacteria possess other mechanisms of action on snRNP’s it may be an interesting focus of future studies.

### Sheep interdigital skin microbiota and scar tissue formation in footrot

The host response to the skin microbiota has to be carefully regulated as innocuous microbes and the host surveillance at epithelial barriers are in constant close proximity. In healthy tissues bacteria are located predominantly in the epidermis while tissue damage and invasive bacteria such as *D. nodosus* allow access of bacteria into deeper dermal tissue layers (10). The ovine host response to footrot demonstrated through differential expression of a range of transcripts involved in proinflammatory mediation (cytokines; IL-19, IL-20, IL-6, LIF, chemokines; CCL2, CCL20, CXCL1, CXCL8 and prostaglandin-endoperoxide synthase 2; PGE2/COX2), of matrix metalloproteases (MMP1, MMP3, MMP9, MMP13, TNC, TIMP1) and interestingly their regulators (SERPINE1, ADAMTS4, ADAMTS16) during footrot, all of which are associated with wound healing, collagen turnover and scar tissue formation. Collagen I was detected significantly more in diseased dermis than in non-infected dermal tissue leading to the conclusion that infection or co-infection clearly indicates current, or ongoing, scar formation in the dermis.

The process of second intention wound healing with scar formation is classically divided into three main overlapping phases: inflammation, proliferation, and remodelling. Localised inflammation is the first response to any breach of haemostasis with the initiation of cytokine and chemokine production leading to neutrophil and macrophage recruitment to the site of inflammation (31). The cytokines and chemokines IL-6, CCL2, CXCL1, CXCL8 identified as differentially expressed in response to footrot, are associated with acute inflammation in response to tissue injury (31, 32). In normal skin wound healing, the inflammation usually lasts for 2–5 days and ceases once the harmful stimuli have been removed. The IL-20 cytokine family (IL-19, IL-20, IL-22, IL-24, IL-26) contribute to various stages of this wound healing process: they are primarily secreted by infiltrating innate immune cells and lymphocytes shortly after an injury. Initially released by infiltrating macrophages, they preferentially stimulate keratinocytes to secrete antimicrobial peptides and chemokines, in order to reduce infection and accelerate inflammation, and to produce increased levels of vascular endothelial growth factor A (VEGFA), which in turn promotes angiogenesis. IL-20 subfamily cytokines directly stimulate keratinocyte proliferation and migration, and indirectly support the proliferation of keratinocytes by enhancing the production of epidermal growth factor (EGF) and keratinocyte growth factor (KGF) (33). Surprisingly, we observed significantly reduced expressed IL-19 and IL-20 transcripts in footrot samples. This was accompanied by reduced expression of secretory leucocyte protease inhibitor 1 (SLP1), a protein essential for optimal wound healing due to its antimicrobial and anti-inflammatory properties (34). Macrophages, initially producing pro-inflammatory mediators, transition in response to local immune signals to an anti-inflammatory phenotype. This promotes the resolution of inflammation and a transition to the proliferation phase of second intentiaon wound healing focusing on re-epithelialisation through migration and proliferations of keratinocytes, deposition of type III collagen, and angiogenesis (31). Matrix metalloproteases (MMPs) are crucial to this phase as extracellular matrix degradation and deposition is essential for wound re-epithelialisation and also during tissue remodelling. MMP expression and activity are tightly controlled during wound healing, at the expression levels and through endogenous tissue inhibitors of metallo proteases (TIMPs); specific MMPs are confined to particular locations in the wound and to specific stages of wound repair (35). MMP-1, MMP-3, and MMP-9 are the major chemokine regulators during wound healing, degrading chemokines by proteolysis and promoting the transition to the proliferation phase. MMPs-1, 8, 9 and 13 are transiently upregulated to remodel the fibrin clot and replacing it with new extracellular matrix. In addition, they are fundamentally stimulating the migration of keratinocytes into the wound bed. During re-epithelialisation, keratinocytes migrate from the surrounding epithelium and proliferate to achieve wound closure. This is accompanied by a decreased expression of pro-migratory MMPs (MMP-1 & 2) and an increased tissue remodelling MMP-3 expression (35, 36).

The dysregulation of MMPs leads to prolonged inflammation and delayed wound healing (37). In footrot affected tissues we observed differential expression of the collagenases MMP-1 and 13, the gelatinase MMP-9 and the stromalysin MMP-3. It is well established that high levels of MMP-1 lead to defective re-epithelialisation, with MMP-13, expressed deeper in the tissues, leading to granulation tissue formation (37). Increased MMP-9 levels are consistent with chronic wounds, leading to the reduced expression of the growth factors required for the healing process while prolonging the inflammatory phase (37). The stromalysin MMP-3 is expressed by proliferating keratinocytes at the distal end of the wound and is essential for wound healing (37). However, we observed reduced expression compared to uninfected interdigital skin tissue, which would impact on the ability of infected interdigital skin tissue to heal. Long chain fatty acids and collagens are essential for skin barrier function (36, 38). The increase in expression of fatty acid elongases (ELOVL]7, ELOVL3, ACSBG1) and of proteins involved in collagen production and collagen binding in response to footrot suggests some level of skin regeneration is ongoing.

One of the bacteria significantly increased in abundance on footrot-infected lesions, *M fermentans*, might affect the ability of host skin cells to respond to bacterial infection. Chronic infections of monocytes and macrophages with intracellular low pathogenic Mycoplasma spp. of have been shown to impair their inflammatory response to live bacteria and bacterial products (39, 40). That we see higher transcript levels of outwardly healthy interdigital skin is in contrast to the lack of detection of MMP RNA in healthy human or murine skin (41). However, this is consistent with a marked expression of the inflammatory cytokines/chemokine IL1β, IL6 and CXCL8 in outwardly healthy ovine interdigital skin, which might be due to the constant environmental changes and pressures impacting on interdigital skin or might be associated with subclinical disease that may have developed into ID and footrot in the future. During the remodelling phase of scar tissue formation, initially deposited collagen-III molecules are gradually replaced by type I collagen and their orientation becomes more organised (36). Mature cutaneous scars consist of 80-90% type I collagen arranged in parallel bundles (42). This particular orientation as well as less pronounced or missing rete ridges weaken the strength of the scar tissue compared to normal skin in humans to only 70-80% (36). This renders the tissue more susceptible to injury and trauma which are suspected predisposing factors of footrot. The latter might also contribute to the frequently observed relapses and underlines the not only polymicrobial but rather multifactorial aetiology of footrot. For BDD a dysfunctional skin barrier and disturbed tissue integrity is hypothesised to be an essential prerequisite for infection altogether since experimental disease models without skin maceration prior to infection fail to mirror naturally occurring BDD lesions appropriately (43).

Currently there is conflicting evidence of the impact of the microbiome on wound healing, with some evidence of host commensal interactions promoting wound healing while colonisation of pathogenic bacteria may invade deeper into tissues or lead to chronic infections and biofilm formation (44, 45). In the context of footrot, we identified another bacterial species in addition to *D. nodosus* that is associated with footrot and also known to be a synergistic wound pathogen, *P. asaccharolytica*. When present in combination with anaerobic and aerobic bacteria such as *Prevotella melaninogenicus*, *Peptostreptococcus micro* and *Klebsiella pneumoniae, P. asacharolytica* exacerbates the disease process (46–48). While antibiotic injections will affect indiscriminately on commensal and pathogenic bacteria, the effectiveness of parenteral antibiotics in footrot demonstrate their high impact on the pathogenic bacteria leading to swift recovery in most cases (49). Interestingly, resistance genes against tetracycline, the most commonly used antibiotic against footrot have so far not been identified in *D. nodosus* genome sequences, suggesting that the antibiotic treatment mainly affects other microbes of that polymicrobial infection enabling host immune system to eliminate *D. nodosus*.

## Conclusion

Ovine footrot is a complex polymicrobial disease and there is a clear need to further elucidate the intricate host microbial interactions. We aimed to investigate the host response as well as the microbial taxa in tissues and their intra-tissue expression levels using metatranscriptomics in naturally infected tissues. It is well published that skin damage is required to allow *D. nodosus* infection to establish (2, 50). As expected, the host response in footrot is characterized by differential expression of proteins with roles in wound healing and chronic wounds. As in the absence of *D. nodosus*, interdigital dermatitis resolves, the presence of *D. nodosus* may be essential to allow the establishment of the microbes associated with under running footrot, including *P. asaccharolytica*. In these later stages of disease, the presence of those bacteria, such as *M. fermentans*, may contribute to a dampening of the immune response unable to remove the invading bacterial pathogens leading to chronic infection.

## Materials and Methods

### Sample collection

Sheep were assessed post slaughter for foot health. Any individual animals showing signs of footrot were selected for sample collection. Debris was removed from all the feet and cleaned using purified water. Sterile nylon flock swabs (E-swabs 480CE, Copan U.S.A.) were taken from the interdigital space and stored in liquid Amies media at 5°C overnight. The foot was then washed with a chlorohexidine solution (National Veterinary Services, U.K.). Any hair was removed from the feet with scissors, prior to the collection of an 8mm biopsy using a punch (National Veterinary Services, U.K.). Biopsies were placed in RNALater (Sigma Aldrich, U.K.), stored at 5°C overnight before being frozen at −80°C.

### DNA Extraction from swabs

The interdigital space swabs were placed on a MixMate (ThermoFisher, U.K.) for 5 minutes at 800rpm to thoroughly disperse the bacteria in the amies solution from the swab. The liquid was transferred into a low-bind 1.5ml tube and centrifuged at 12,000 rpm for 5 minutes. The supernatant was removed, and the pellets were resuspended in 200μl of RNAse-free molecular biology grade water (Thermo Fisher, U.K.) (51). DNA was isolated using the Qiagen Cador Pathogen Mini Kit, following the manufacturer’s guidelines, eluted in 60μl of elution buffer. The DNA samples were quantified using the Qubit 3.0 and dsDNA high sensitivity dye (Qiagen).

### RNA Extraction

Biopsies were thawed on ice before being cut into approximately 30mg sections. One section was added to a MACs M tube (Miltenyi Biotech, U.K.) containing 1ml Qiazol (Qiagen, U.K.) and dissociated on a GentleMACs (Miltenyi Biotech, U.K.) using the manufacturers RNA settings. The sample was centrifuged and incubated at room temperature (RT) for 5 minutes before transferring the lysate to a 1.5ml centrifuge tube. Proteinase K (20μl) was added to the sample before being incubated at 56°C for an hour. Chloroform (200μl) was added and shaken vigorously for 15 seconds. The sample was then incubated at RT for 2 minutes before being centrifuged at 12,000xg for 15 minutes at 4°C. The upper aqueous phase was transferred to a fresh 1.5ml centrifuge tube before the addition of 1x volume of 70% ethanol. The sample (up to 700μl) was added to an RNeasy Mini Spin column (Qiagen, U.K.) and centrifuged at RT at 8000xg for 30 seconds. Any remaining sample was also passed through the column. All remaining steps followed the manufactures guidelines with elution in 30μl of RNAse-free molecular biology grade water (Thermo Fisher, U.K.).

### Dual RNA Sequencing

The extracted RNA was quantified using the Agilent Bioanalyser RNA Nano 6000 kit. Healthy foot sample RNA with a RIN score of ≥7 and footrot sample RNA with a DV200 >85 was chosen for sequencing. The samples were treated using the RiboZero Gold (epidemiology) ribosomal depletion kit (Illumina, U.S.A.) and prepared for sequencing using Illumina TruSeq library preparation (Illumina, U.S.A.). The samples were sequenced on a HiSeq 3000 using 150bp paired end chemistry (Leeds Institute of Molecular Medicine, sequencing facility) at 6 libraries per lane over 14 lanes giving approximately 75 million reads per sample.

### Data Analysis

All analysis was carried out using default settings unless stated. Raw reads were analysed for quality and adaptor removal using Skewer (52). An initial step for the RNASeq data consisted of aligning the reads with HISAT2 (53) against the sheep genome (Oar_v3.1, downloaded 21/07/2017) (54) to separate the ovine and potential bacterial transcripts. The sheep reads were then parsed for transcript alignment using Kallisto (55) and the sheep genome (Oar_v3.1, downloaded 21/07/2017) (54). Differential analysis was calculated with Sleuth (56). The differentially expressed genes were imported in R (57) and clustered using BiClust and the Block Correlated Coupled Clustering (58). The reads which did not align to the sheep genome and the metagenomic reads were used as input for taxonomic assignment and bacterial populations were determined with Kraken (59), false positives were identified with KrakenUniq (60) and results were filtered with MAG_TaxaAssigner (https://github.com/shekas3/BinTaxaAssigner).

### Confirmation of selected bacterial species by PCR or qPCR

Parallel tissue samples from the same foot as for RNA isolation were used. Tissue homogenisation and DNA extractions were performed as described previously using QIAamp Cador kit (Qiagen) (61). The DNA samples were quantified using the Qubit 3.0 and dsDNA high sensitivity dye (Qiagen). Bacterial load was quantified using real-time PCR based on 16S rRNA gene for eubacteria (62) and *D. nodosus* (63), *F. necrophorum* subspecies necrophorum primers targeted the gyrB gene (51). *M. fermentans* sequences were amplified by species specific PCR for 16S rRNA (64) and *P. asaccharolytica* sequences were amplified by PCR (65) followed by sequencing (Eurofins Genomics).

### Correlation Testing

Both tests were performed using the same dataset. For each of the samples (biopsy, swab and host), each of the bacteria species were labelled as either present or absent in each sample. A difference score was prescribed if species was present in one of the swab or biopsy samples from a sheep but, not both (i.e., the swab and biopsy gave different results for the presence of the species). If present in both or neither the biopsy nor swab samples for then. A t-test statistic was calculated by taking the sum of all differences across all sheep. To perform the randomisation procedure the labels of the original 52 samples (sheep and sample type) were randomly reassigned and the test statistic recalculated. This procedure was repeated 10 000 times, giving randomised test statistics.

### Host Pathogen Interactions

To identify putative host pathogen interactions the correlation script PHInder was used (https://github.com/addyblanch/PHInder). Briefly, samples with zero assignments were removed, and a minimum presence value was set to one. A new data matrix was formed and the hypergenometric distribution was calculated using phyper (https://www.rdocumentation.org/packages/stats/versions/3.6.2/topics/Hypergeometric) and a probability, producing a significance (p) value and adjusted p value.

### Tissue staining, image capture and analysis

Biopsies were processed in ethanol and xylene, mounted in paraffin and 7 µm thick serial sections collected throughout the tissue onto polysilinated microscope slides. The paraffin from each tissue section was melted at 60ᵒC for 5-10min, followed by immersed in xylene twice for 5min each to remove the paraffin. Tissue sections were then rehydrated in 100% ethanol, 90% ethanol, 70 % ethanol and distilled water for 5min each. Picrosirius red (PS) stain was used to differentiate and quantify collagen types I, III and undifferentiated collagens using the picrosirius stain kit (Polysciences, Inc., Pennsylvania, USA). The observer was blinded to the sample identification to avoid subconscious bias. Images were captured using a Leica CTR500 microscope (Leica Microsystems, Germany) with or without polarised light. For each sample, three sections approximately 400 µm apart were analysed. At 40x magnification, five non-overlapping photos were taken from each section from the dermis of PS stained sections (330 photomicrographs analysed).

For 13 healthy and 8 footrot samples, 15 photomicrographs per sample were captured (systematic random sampling) and analysed using Image-Pro Plus (Media Cybernetics, Inc, Pennsylvania, USA) to quantify the area of collagen in each image, with separate measurements for type III (green), type I (yellow) and undifferentiated (red) collagen (315 images analysed in total). Total collagen proportion was calculated as the sum of type III, type I and undifferentiated collagen proportions.

### Statistical analysis

The taxonomic count data were analysed for statistically significant differences in R (57) using the edgeR wrapper (66) as part of the Phyloseq package (67). Diversity statistics were calculated using vegan (68) and differences were calculated using Mann-Whitney U tests in Prism 8.01 (GraphPad Software Inc. USA).

Statistical analyses of histology images were performed on GraphPad Prism version 6 for windows. Resulting data were presented as frequencies and percentages and were analysed by student T-test or Kruskal Wallis test, dependent on data distribution. Analysis was taken as significant when p ≤ 0.05.

## Author Contributions

AMB and ST developed the idea, designed, and supervised the experiments and alongside SW, CMB, JKM and GE have written the manuscript. AMB completed the bioinformatics analysis. CES, CMB and JKM carried out the lab work, NN performed and analysed bacterial qPCR. CR and CB performed and analysed the collagen assays. LS performed additional statistical and mathematical assessment of the data. All authors have read the manuscript.

## Ethical Statement

This study was reviewed and approved by the University of Nottingham, School of Veterinary Medicine and Science ethical review committee ERN: 1144 140506 (Non ASPA).

## Acknowledgements

The authors would like to thank Professor Christoph Mülling for helping facilitate this research collaboration with the Faculty of Veterinary Medicine, Leipzig University. The authors would also like to thank Dr Ian Carr and Ummey Hany at The Leeds Institute for Molecular Medicine Sequencing Facility for all their experimental suggestions and sequencing support.

## Open Access Data

All sequence data generated for this study is held in the NCBI SRA under the accession number PRJNA725378.

